# Structure basis of the phytoplasma effector SAP05 recognition specificities to plant Rpn10 in ubiquitin-independent protein degradation

**DOI:** 10.1101/2023.08.24.554544

**Authors:** Liying Zhang, Yunxiang Du, Qingyun Zheng

**Author notes:** These authors contributed equally to the work (Liying Zhang, Yunxiang Du). Author contributions: L.Z., Y.D., Q.Z. designed research; L.Z., Y.D. performed research; all authors wrote and edited the paper. This open access article is distributed under Creative Commons Attribution License 4.0 (CC BY).

## Abstract

Recent research reported an effector protein SAP05 secreted by phytoplasma could hijack hostage Rpn10 subunit of proteasome and target degradation of GATA or SPL transcription factors (TFs) without ubiquitin. As a result, plant phenotypes will be reprogrammed to a state suitable for phytoplasma amplification and insect vector propagation which present prolonged growth cycles, witches’ broom-like proliferations of leaves and sterile shoots. We are curious about how could SAP05 target degradation bypassing the ubiquitin pathway and if there is any conserved interaction between different species of SAP05 with AtRpn10. Here, we have determined the crystal structure of the AtRpn10 complex with AY-WB or OY SAP05. Structure alignment revealed similar recognition patterns through sequences of two species SAP05 are poor homologous, which suggested phytoplasma may conduct resemble mode in plant infection. After docking the complex structure to the plant proteasome, we found that SAP05’s location is near the ATPase central pore, which may be close enough to submit substrate with no need for ubiquitin in the degradation process.

## Introduction

The ubiquitin proteasome system (UPS) is a major pathway for protein quality control in eukaryotic cells. It relies on a cascade enzyme system consisting of E1s-E2s-E3s to covalently modify ubiquitin (Ub) on substrate proteins as a degradation tag, selectively degrading misfolded or damaged proteins to prevent their accumulation^1^. Dysregulation of the UPS is associated with a wide range of human diseases, such as cancers and degenerative diseases^2^. A variety of enzymes in the pathway are therapeutic targets including the proteasome. Moreover, hijacking the endogenous UPS to degrade disease-related proteins has become a therapeutic approach of great interest in the biomedical field, such as the development of small molecules like molecular gels and PROTACs^3^. These synthetic molecules could establish contact between E3 ligase and disease proteins, and mediate the generation of ubiquitin chains on target proteins with the help of E3 ligase to make the target proteins recognized by proteasome for degradation.

Interestingly, a subset of eukaryotic proteins have been shown to undergo ubiquitin-independent proteasomal degradation (UbInPD), including ornithine decarboxylase, p53 and its downstream apoptosis regulators BIM_EL_, NPK as well as cyclin-dependent kinase inhibitor p21^4-7^. These proteins usually contain an unstructured region, which can be recognized and degraded by the proteasome in *vivo* and in *vitro*, despite that all Lys sites are blocked to ubiquitination. Recently, global protein stability (GPS)-peptidome technology has been applied to identify novel full-length protein substrates that are degraded without ubiquitin^8^. They identified thousands of UbInPD substrates, some of which required C-terminal degradation sequence (C-degron) and shuttling factors of the Ubiquilin family for proteasome recognition. It suggests the process that UbInPD regulates substrate turnover in vivo is more common than previously appreciated.

In addition to regulating eukaryotic homeostasis, pathogen-encoded effector proteins also can hijack some hostage proteins for direct proteasomal degradation through ubiquitin-independent pathways. An insect-vectored parasitic phytoplasma AY-WB (Aster Yellows phytoplasma strain Witches’ Broom) infects the plant host Arabidopsis thaliana and secretes the effector protein SAP05^9^, which induces almost all extended developmental phenotypes of the phytoplasma, such as plant dwarfism, prolonged growth cycle, increased number of leaves and lateral branches, and infertility of flowers, etc. A recent study found that SAP05 proteins bind the host Rpn10 in the 26S proteasome and plant GATA or SPL transcription factors, which results in target degradation without ubiquitination. Therefore, concurrent destabilization of SPLs and GATAs by SAP05 effectors decouples plant developmental transitions. Detailed assays in vivo and in vitro demonstrated that SAP05 forms a bridge between the SPL5 ZnF domain and Rpn10 vWA domain to generate a ternary complex, and ubiquitination of lysine residues within SPL5 may not be required for SAP05-mediated degradation. Besides AY-WB SAP05, some homologs have evolved to interact and degrade other families of transcription factors in plant MADS-domain transcription factor (MTFs)^10-13^, while SAP05-induced Rpn10 interaction is not conserved in *homo sapiens*. This distinct mode of action given that targeted protein degradation (TPD) has been applied to engineer plant Rpn10 conferring resistance to SAP05. Due to the wide range of phytoplasma targets and the strong spreading ability assisted by insect hosts, which seriously affects crop yields including grain and causes incalculable economic losses, obtaining structural information about the molecular mechanism of phytoplasma effector protein hostage proteasome-targeted degradation will be helpful to more accurately guide the transgenic operation of crop pest resistance, and also facilitate the development of novel chemical inducers of degradation (CIDEs) that directly target proteasome^14^.

Here we resolved the crystal structure of phytoplasma AY-WB_SAP05 protein in complex with Arabidopsis thaliana Rpn10vWA domain. The structure reveals that AY-WB_SAP05 binds to AtRpn10 predominantly through a V-shape peptide as well as a C-terminal peptide which constitutes the surface with hydrogen-bond linkages, polar contacts, and hydrophobic interactions. This recognition pattern is highly consistent with the recognition of Arabidopsis Rpn10 by the poor sequence homologous onion yellow phytoplasma (OY-M strain) SAP05, as indicated by the crystal structure of the complex of OY_SAP05 and Rpn10vWA. The resemblance in structure suggests that phytoplasmas may have a similar mode to infect plants by attaching proteasomal subunits to destabilize development-related transcription factors, as a result affecting plant phenotypes. To understand how SAP05 interacts with Rpn10 at the proteasome level, we docked the structure of the SAP05-Rpn10 complex with the Spinach proteasome and found that there was no significant change in the relative structure of Rpn10vWA. Therefore, we hypothesized that the SAP05 protein may play a molecular glue-like role in mediating the direct degradation of transcription factors by bringing transcription factors closer to the ATPase central pore of proteasomes without undergoing an additional ubiquitination process.

### Phytoplasma effector SAP05 recognizes proteasome subunit Rpn10 in a host species-specific manner

Previous data indicated that aster yellows witches’-broom phytoplasma SAP05 (AYWB_SAP05) mainly recognizes the N-terminal vWA (von Willebrand A) domain of *A. thaliana* Rpn10 (AtRpn10)^9^. We therefore constructed and recombinantly expressed the mature SAP05 without signal peptide sequence SVM (33-135 amino acids, and just called SAP05 for convenience hereinafter) and the GST-tagged AtRpn10 vWA domain. Accordingly, surface plasmon resonance (SPR) experiments showed that the AYWB_SAP05 bound AtRpn10 with three orders of magnitude lower dissociation constant ((0.4 ± 0.039) μM) compared with the *homo sapiens* Rpn10 (hRpn10) vWA (Figure 1A), which is consistent with the previous finding that SAP05 cannot recognize human proteasome. We also tested another onion yellows phytoplasma OY_SAP05 that is homologous to AYWB_SAP05 with a relatively low conserved surface. It also showed a high affinity against AtRpn10 instead of hRpn10, which is similar to AYWB_SAP05 (Figure 1B). These findings suggest a conserved recognition behavior between phytoplasma SAP05 homologs and host proteasome subunit Rpn10, and Rpn10 of *homo sapiens* have evolved to escape this phytoplasma effector recognition.

**Figure 1.**
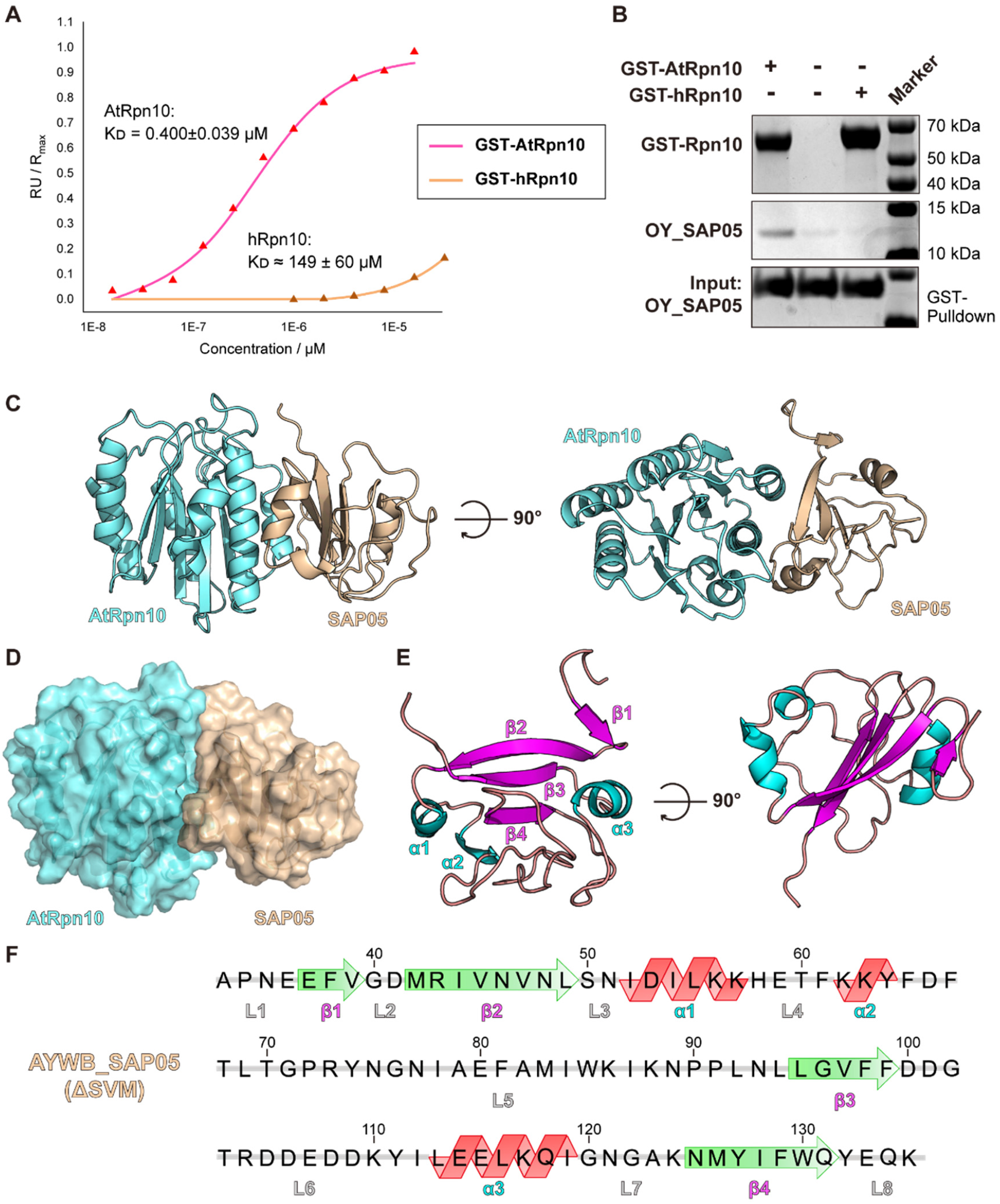
Crystal structure of AYWB_SAP05-AtRpn10 complex. A) The curve of response value versus concentration of GST-tagged AtRpn10/hRpn10 in the SPR binding assay of Rpn10 binding to immobilized AYWB_SAP05. The dissociation constant between GST-hRpn10 and AYWB-SAP05 is far larger than GST-AtRpn10 -AYWB_SAP05 pair. B) GST-pull down of tagged AtRpn10 or hRpn10 with the appearance of analyte OY_SAP05. C) Two views of the cartoon model of AYWB_SAP05-AtRpn10_vWA global structure. D) Surface of the AYWB_SAP05 and AtRpn10 in structure. E) F) Secondary structures and their sequence and number in AYWB_SAP05.

### Structures of AYWB_SAP05-AtRpn10 complex

To gain structural insight into how AYWB_SAP05 hijacks the proteasome subunit Rpn10 in *A. thaliana*, we determined the crystal structure of the AYWB_SAP05-AtRpn10_vWA complex. Recombinant expressed AYWB_SAP05 and AtRpn10 truncates were able to form a stable 1:1 heterodimeric complex in gel filtration analysis (Figure S1A). The purified protein complex of AYWB_SAP05-AtRpn10 was collected and crystals were grown from sitting drop with reservoir buffer containing 0.2 M Sodium malonate, 20% w/v Polyethylene glycol 3,350. Utilizing the molecular replacement with Alphafold2-enabled models of two proteins (Rpn10: AF-P55034-F1, SAP05: AF-Q2NK94-F1), resulting in a final structure at 1.57 Å resolution (Figure 1C and 1D).

The AtRpn10_vWA in our structure adopts a conformation similar to that predicted by Alphafold2, which also resembles the structure of vWA in the spinach 26S proteasome^15^ (PDB 8AMZ, rmsd. = 0.781 Å) (Figure S1B). And the SAP05, which previously had no homologous structure, performs to fold as a globular shape and is identical to the prediction of Alphafold2 except for the smaller amount of β-sheet secondary structure (Figure S1C). The SAP05 structure contains a four-strand β-sheet core layer and three α-helices distributed on either side (Figure 1E and 1F. There are two long and compact loops, L5, L6, between α2 and β3 and between β3 and αC respectively, wrapping together on the opposite side of the SAP05-Rpn10 interface.

### A non-classical interface of AYWB_SAP05 bound to AtRpn10

SAP05 and Rpn10 form a 1783.1 Å^2^ quite extensive interface, containing 6 pairs of hydrogen-bond linkages and a large area of polar contacts and hydrophobic interactions (Figure 2A and 2B). A V-shape peptide (43-61 aa.) in SAP05, which uses L3 (S50, N51) as the vertex, β2 and α1-L4 as two sides (called by β-side and α-side hereinafter respectively), is involved in the AtRpn10 recognition (Figure 2C). This V-shape peptide docks on the α1 and α2 of AtRpn10_vWA. In detail, the vertex S50 side chain, N48 main-chain carbonyl oxygen on the β-side and H58 side chain on the α-side form three hydrogen bonds with AtRpn10 E31, Q27, and N34 respectively (Figure 2A). SAP05 R43 and T60, which are located at the N-termini and C-termini of the V-shape peptide, have polar contacts with AtRpn10. And the V45, V47, and L49 on the β-side and F61 on the α-side of SAP05, contribute to binding by hydrophobic effect simultaneously (Figure 2C).

**Figure 2.**
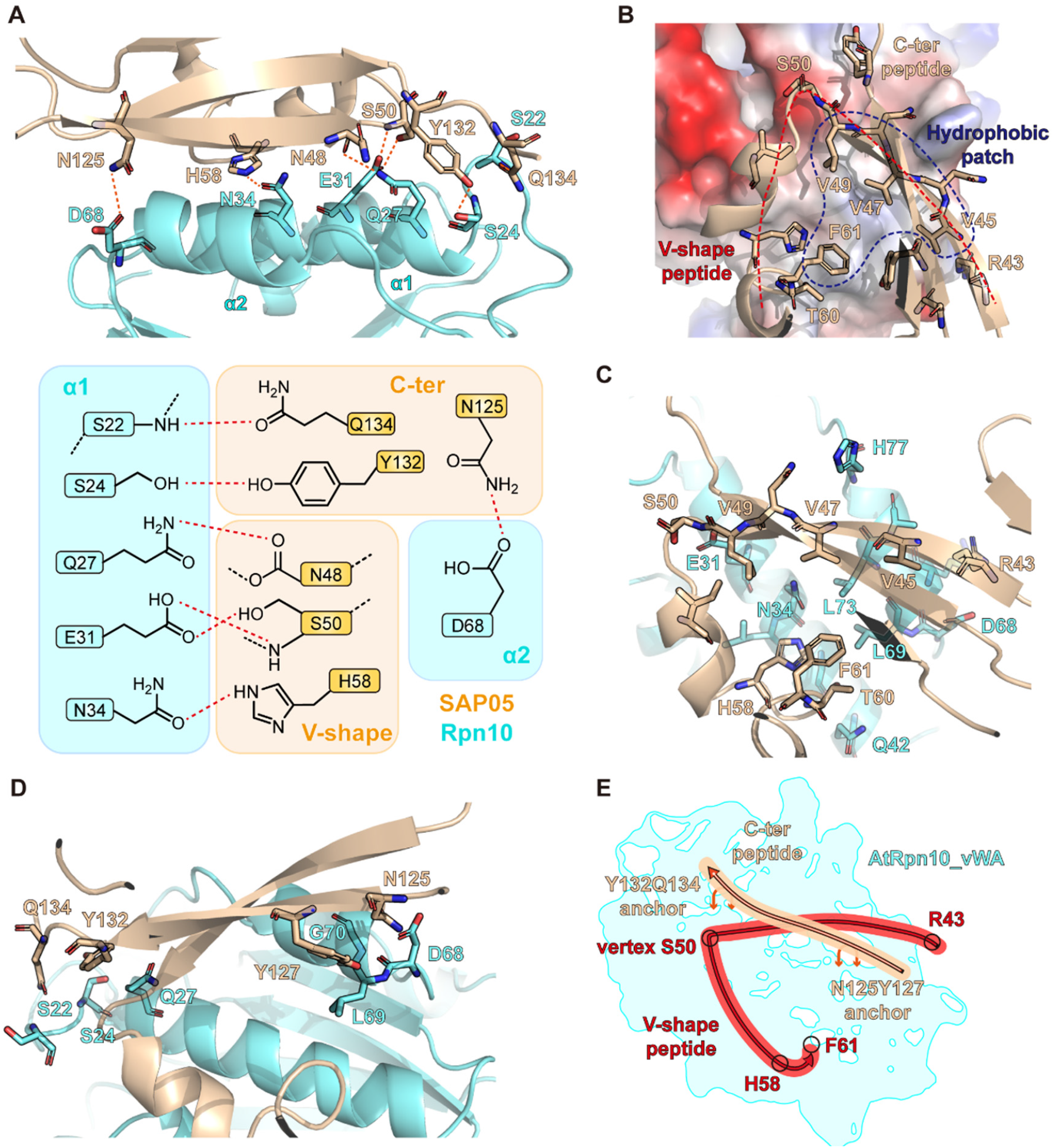
Detailed interaction between AYWB_SAP05 and AtRpn10. A) Hydrogen bonds formed between AYWB_SAP05 and AtRpn10. The amino acids involved in the hydrogen bonds formation are shown as sticks in the upper figure. The lower diagram summarizes the hydrogen-bond interactions between different regions of two proteins. The hydrogen bond was represented by red dashed lines. B) Components of AYWB_SAP05-AtRpn10_vWA interface. The vacuum electrostatic potential surface of AtRpn10 and two key binding regions on SAP05: V-shape peptide and C-ter peptide are shown as the cartoon model. C) Interactions between V-shape peptide and AtRpn10. The amino acids involved in the interaction are shown as sticks. D) Interactions between N-ter peptide and AtRpn10. E) Cartoon diagram of how AYWB_SAP05 interacts with AtRpn10_vWA domain, vWA surface is colored by cyan. V-shape peptide and C-ter peptide are indicated in red and wheat lines respectively.

Meanwhile, C-terminal β4 and L8 (called C-ter peptide as a whole) of SAP05 form an extra two-point anchoring conformation on the surface of AtRpn10, straddling the β2 strand and “stapling” it onto the interface (Figure 2D). On the SAP05 C-termini side, Y132 forms a hydrogen bond with AtRpn10 S24 while Q134 interacts with the main chain of the N-terminal of AtRpn10 L1. On the other side, SAP05 N125 and AtRpn10 D68 also take a hydrogen-bond contact. In summary, SAP05-Rpn10 recognition is mediated by the cooperation of polar contact including a large number of hydrogen bonds, and hydrophobic interactions (Figure 2E).

Proteasome subunit Rpn10 is a well-known “ubiquitin receptor” which comprises the quantity different C-terminal ubiquitin interacting motifs (UIMs) to mediate the direct recognition of ubiquitin, ubiquitin-like (UBL) domain or UBL proteins (ATG8^16^, FAT10^17^). Previously, shuttle factors such as the Ubiquilin family, DSK2, and RAD23 were reported to interact with Rpn10_UIM to transport ubiquitinated substrates or induce the UbInPD directly. And even the vWA of Rpn10 is able to act as an atypic region bound to FAT10 and NUB1L UBL domain. Here, we found Rpn10 vWA domain recognizes the special exogenous shuttle factor AYWB_SAP05 which has no similarity to ubiquitin (*i*.*e*. no UBL domain) through a previously unknown atypical interface. Our structure reveals that this atypical recognition depends on a stapled V-shape peptide forming extensive hydrogen-bonds and hydrophobic interactions with the association of the Rpn10 vWA domain.

### Key residues of SAP05 contributing to the conserved recognition of AtRpn10

To validate the interface of the crystal structure, we introduced single or double Ala mutations to key interface residues of recombinant SAP05 or Rpn10 (Figure S2A and S2B) and tested the binding affinity by the SPR measurement. Replacing the SAP05 residues whose side chains were involved in the hydrogen bonds formation with Ala greatly weakened the interaction, whether the residues of V-shape peptide (S50A, H58A) (Figure 3A and 3B), or C-ter peptide (Y132/Q134A). The binding force also decreased when the T60 was turned to Ala, which diminished the polar contacts between the two proteins. Expectedly, the N48A mutation did not affect the interaction between AYWB_SAP05 and AtRpn10, since N48 uses main-chain carbonyl oxygen rather than side-chain as the hydrogen-bond acceptor (Figure 3A and 3B). Consistently, mutations of the key residues of interaction on AtRpn10 also abrogated the SAP05 binding (Figure 3C). Altogether, the residues we observed at the interface are indeed involved in the interaction between AYWB_SAP05 and AtRpn10.

**Fig 3.**
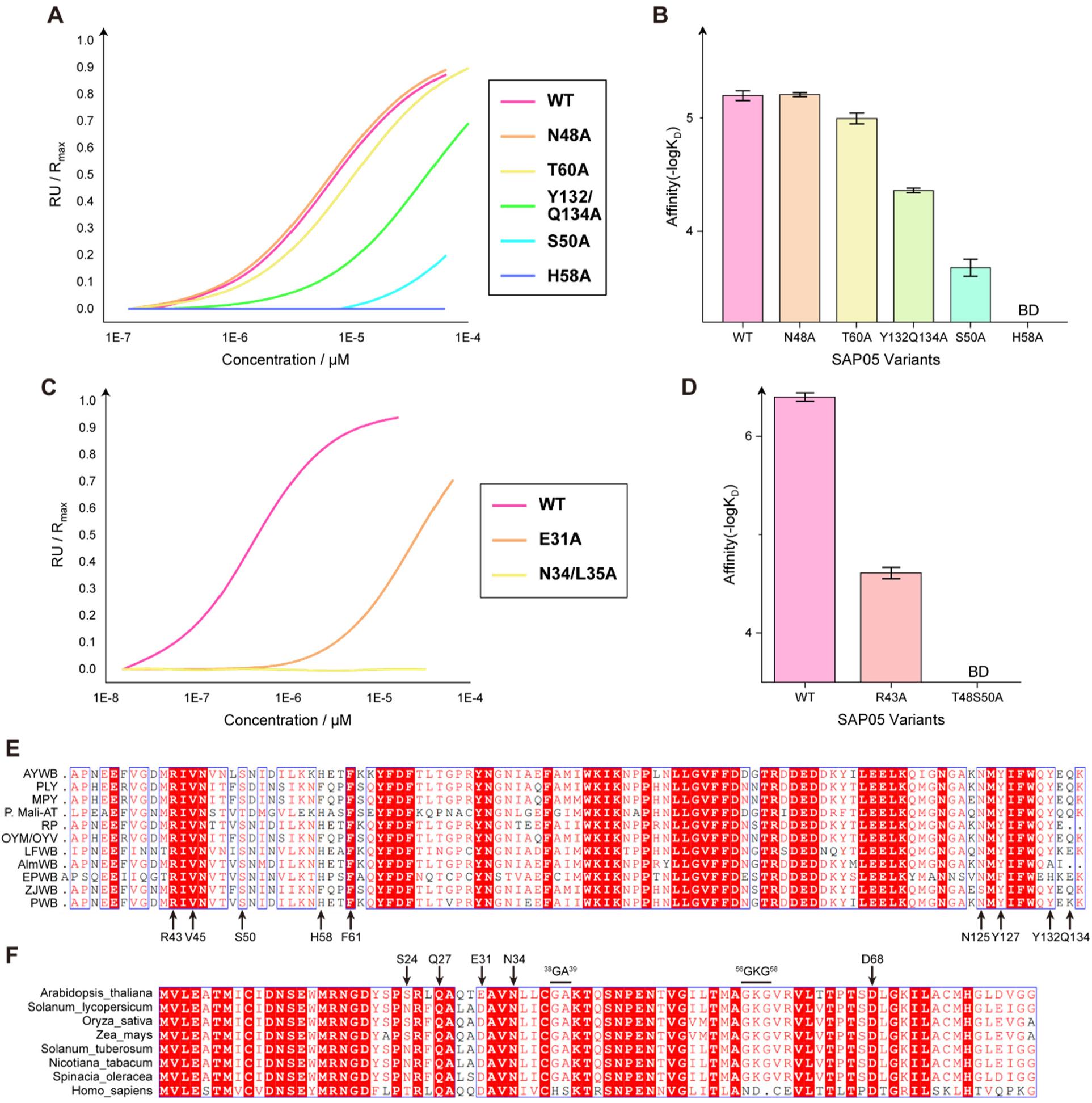
Conserved sequence and binding mode of SAP05-Rpn10 recognition of different species. A) The curve of response value versus concentration of AYWB_SAP05 mutants in the SPR binding assay of SAP05 proteins binding to immobilized AtRpn10. B) The histogram of the dissociation constant between WT AtRpn10 and different AYWB_SAP05 mutants measured by SPR. C) The curve of response value versus concentration of GST-AtRpn10 mutants in the SPR binding assay of Rpn10 proteins binding to immobilized AYWB_SAP05. D) The histogram of the dissociation constant between WT AYWB_SAP05 and different AtRpn10 mutants. E) Multiple sequence alignment of SAP05 homologs from different phytoplasma species. Amino acids which are involved in interaction and highly conserved are pointed with arrows at the bottom. F) Multiple sequence alignment of Rpn10 homologs from different plant species. Amino acids which are involved in interaction and highly conserved are pointed with arrows on the top. Two key mutation sites that plant Rpn10 distinct from the mammalian Rpn10 (PSMD4) are also shown on the top.

We next attended to investigate the conservation of the interaction between SAP05 and proteasomal Rpn10. Comparing the sequence of SAP05 in various phytoplasma species, R43, V45 on the β-side, S50 at the vertex, H58 on the α-side, and N125, Y127, Y132, Q134 on C-termini peptide of SAP05 are highly conserved involving in the interaction with AYWB_SAP05-AtRpn10 (Figure 3D). Most of these residues participate in the hydrogen bond formation. On the other hand, residue N34, D68, and S24 of AtRpn10 corresponding to itself binding partner of H58, N125, and Y132 of SAP05, respectively, are also conserved among different plant species (Figure 3E). Especially, E31 of AtRpn10 is replaced by Asp in most other plant species, which may mediate distinct polar contact to stabilize the V-shape peptide anchoring (Figure 3E). These sequence alignment results indicate that most SAP05 homologs may use similar contacts to interact with plant Rpn10.

### Structural conservation of SAP05 homologs binding to AtRpn10

To further illustrate the conservation of recognizing behavior, we co-crystallized the onion yellows phytoplasma (OY-M strain) SAP05 with AtRpn10_vWA. The V-shape region of OY_SAP05 has only 42.11% identity against the AYWB_SAP05 interface while showing similar binding affinity (Figure 1B). Especially, some vital residues whose mutations lead to the obstacle of SAP05-Rpn10 interaction like H58 and T60, are not conserved in OY_SAP05. We finally resolved the 1.78 Å structure of the AtRpn10_vWA-OY_SAP05 1:1 stoichiometric complex (Figure 4B). Compared to the AYWB_SAP05-AtRpn10 complex structure, both SAP05 and vWA in this homologous structure adopt identical conformations, except that OY_SAP05 forms a triangular β-sheet core layer harboring 6 β-strands, which is larger than AYWB_SAP05 (Figure S3A and S3B).

**Fig 4.**
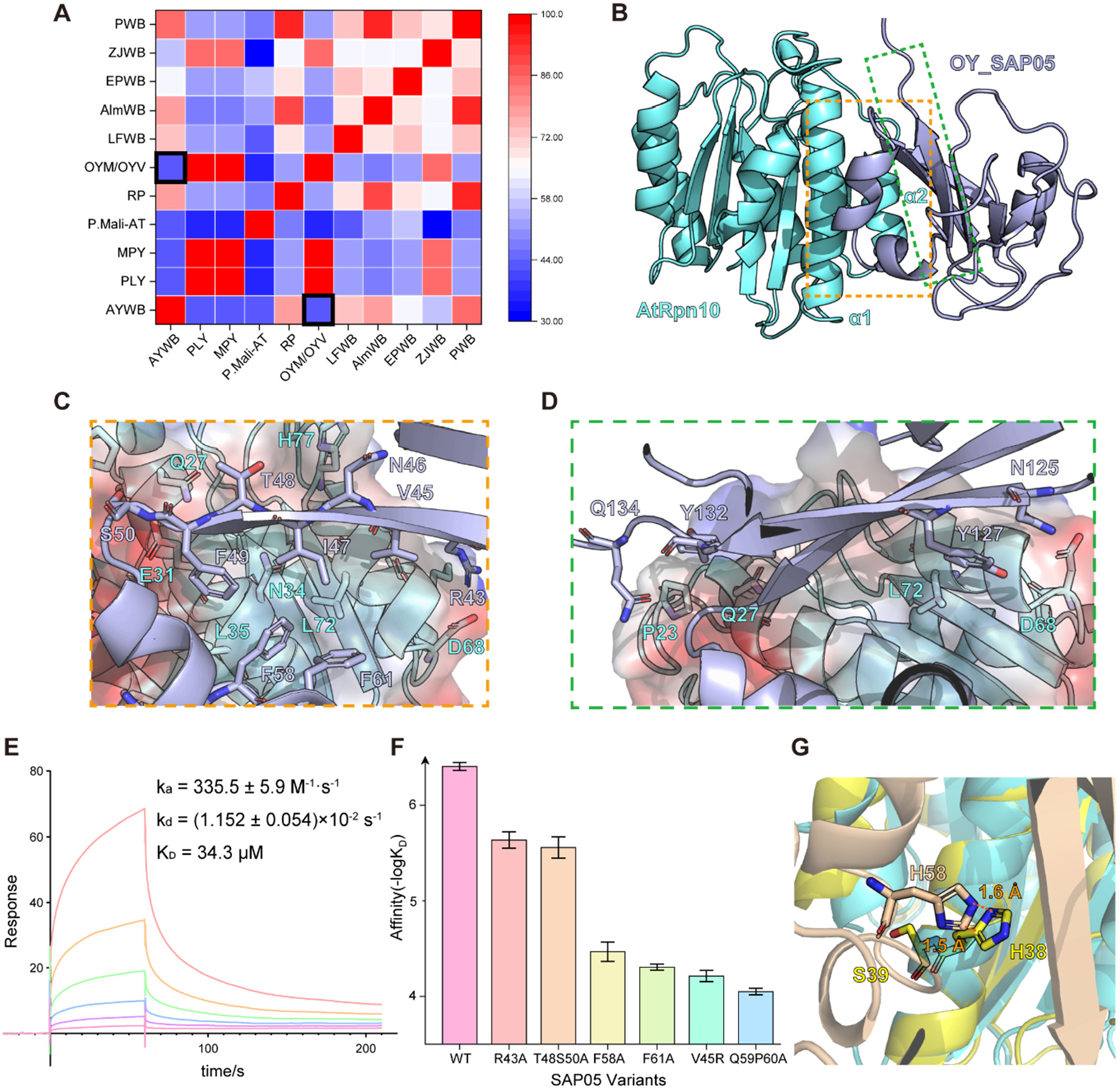
Conserved sequence and binding mode of SAP05-Rpn10 recognition of different species. A) Percent identity matrix of V-shape region of SAP05 homologs in different phytoplasma species. The blocks representing the similarity between AYWB and OYM SAP05 are highlighted. B) Cartoon model of OY_SAP05-AtRpn10_vWA complex global structure. C) Interactions between OY_SAP05 V-shape peptide and AtRpn10. D) Interactions between OY_SAP05 N-ter peptide and AtRpn10. E) Response curve of V-shape-GSGS-C-ter peptide binding to AtRpn10 measured by SPR. F) The histogram of the dissociation constant between WT AtRpn10 and different OY_SAP05 mutants measured by SPR. G) Alignment of our crystal structure to the human Rpn10 (PSMD4 in PDB 6MSB) vWA domain structure. The clash between aligned SAP05 with human Rpn10 was shown in sticks.

Although the sequence similarity between AYWB_SAP05 and OY_SAP05 is relatively low, their recognition manners remain conserved (Figure 4B): The OY_SAP05 docks above the α1 and α2 of Rpn10_vWA. A V-shape peptide (Figure 4C, 43-61 aa.) cooperating with the two-point (N125/Y127 & Y132) anchoring the C-ter peptide (Figure 4D) contributes to the interaction. The vertex of the V-shape peptide contains the conserved S50. Similar to the AYWB_SAP05 structure, the S50, and T48 near the vertex form two hydrogen bonds with AtRpn10 E31, Q27, anchoring the peptide onto the vWA surface (Figure S3C), despite differences in amino acid pairing. The V45, I47, and F49 on the β-side and F61 on the α-side of SAP05 still have hydrophobic interactions with AtRpn10 (Figure S3D). And the conserved R43, which is involved in polar contacts in the AYWB_SAP05 structure, is linked to D68 directly by a hydrogen bond (Figure S3E). Phe-mediated strong hydrophobic interaction covers the contribution of the hydrogen bond involving key residue H58 in the AYWB_SAP05 interface (Figure S3F). In summary, the recognizing conformation of SAP05 remains conserved despite the low sequence similarity. We totally synthesized the fused V-shape and C-ter peptide, which were linked by the GSGS linker. Unfolded peptide appeared to almost completely lose the affinity targeting AtRpn10. After refolded by dialysis and size-exclusion chromatography, the peptide can bind to the AtRpn10 with a K_D_ value of ∼34.3 μM (Figure 4E), which is still far below the binding affinity between the whole AYWB_SAP05 and AtRpn10. Therefore, unlike some sequential specific degron, conformation recognition is necessary for SAP05 to transport transcription factors targeting proteasomal degradation.

The authenticity of this homologous structure is also verified by SPR binding assay involving WT and mutated SAP05. The k_D_ value of WT OY_SAP05 binding with Rpn10 vWA domain is (0.398±0.039) μM, while the affinity of R43, V45, T48S50, F58, Q59P60, F61 mutated SAP05 binding to Rpn10 is reduced by one or two orders of magnitude (Figure 4F). Notably, we failed to prepare V45, F61 mutated recombinant AYWB_SAP05 due to the formation of aggregates. Here, we successfully obtained the corresponding mutants of OY_SAP05, and their performance proved the important role of hydrophobic interaction in the SAP05-Rpn10 recognition. Our data indicated that the phytoplasma SAP05 family may adopt highly similar conformations for recognizing plant Rpn10.

### Changes in interface abrogate the binding of SAP05 to hRpn10

In 2021, Huang et al. reported that AWYB_SAP05 cannot bind with *homo sapiens* Rpn10 homolog PSMD4^9^. They found two key polymorphistic sites in Rpn10/PSMD4 sequence that prevent hsRpn10 from interacting with SAP05: ^38^GA^39^ (*A. thaliana*) to ^38^HS^39^ (*H. sapiens*), and ^56^GKG^58^ (*A. thaliana*) to ^56^ND^57^ (*H. sapiens*). Superposing hsRpn10_vWA (from PDB: 6MSB) to the AtRpn10_vWA in our structure shows that the hsRpn10 ^38^HS^39^ mutation clashes with the SAP05 H58, thus obstructing the SAP05 recognition (Figure 4G). ^56^GKG^58^ to ^56^ND^57^ mutation may prevent SAP05 binding by abolishing the β3 secondary structure and subsequently skewing the important interface α2 (Figure S3G). Our data are consistent with the previous finding and provide clues for the species selectivity of this hijacking process.

### Superposing the SAP05 complex into the proteasome

Rpn10 is located at the periphery of the 19S regulatory particle, where it is in contact with lid subunits Rpn8 and Rpn9 and maintains structural integrity. The α1/α2 surface of Rpn10_vWA does not interact with the other 19S subunit and is accessible for SAP05 binding. By aligning our AYWB_SAP05-AtRpn10 structure with the Rpn10 in the structure of spinach (*Spinacia oleracea*)^15^ proteasome 19S regulatory particle reported recently, we can observe that the SAP05 is placed proximal to the central pore of 19S ATPase OB-ring and has no clash with any constitutive subunit (Figure 5A and 5B). This binding pattern has little impact on the constitutive assembly of proteasome and recruits the transcription factor GATA or SPL to the entrance of ATPase hexamer for the sequential substrate deposition.

**Fig 5.**
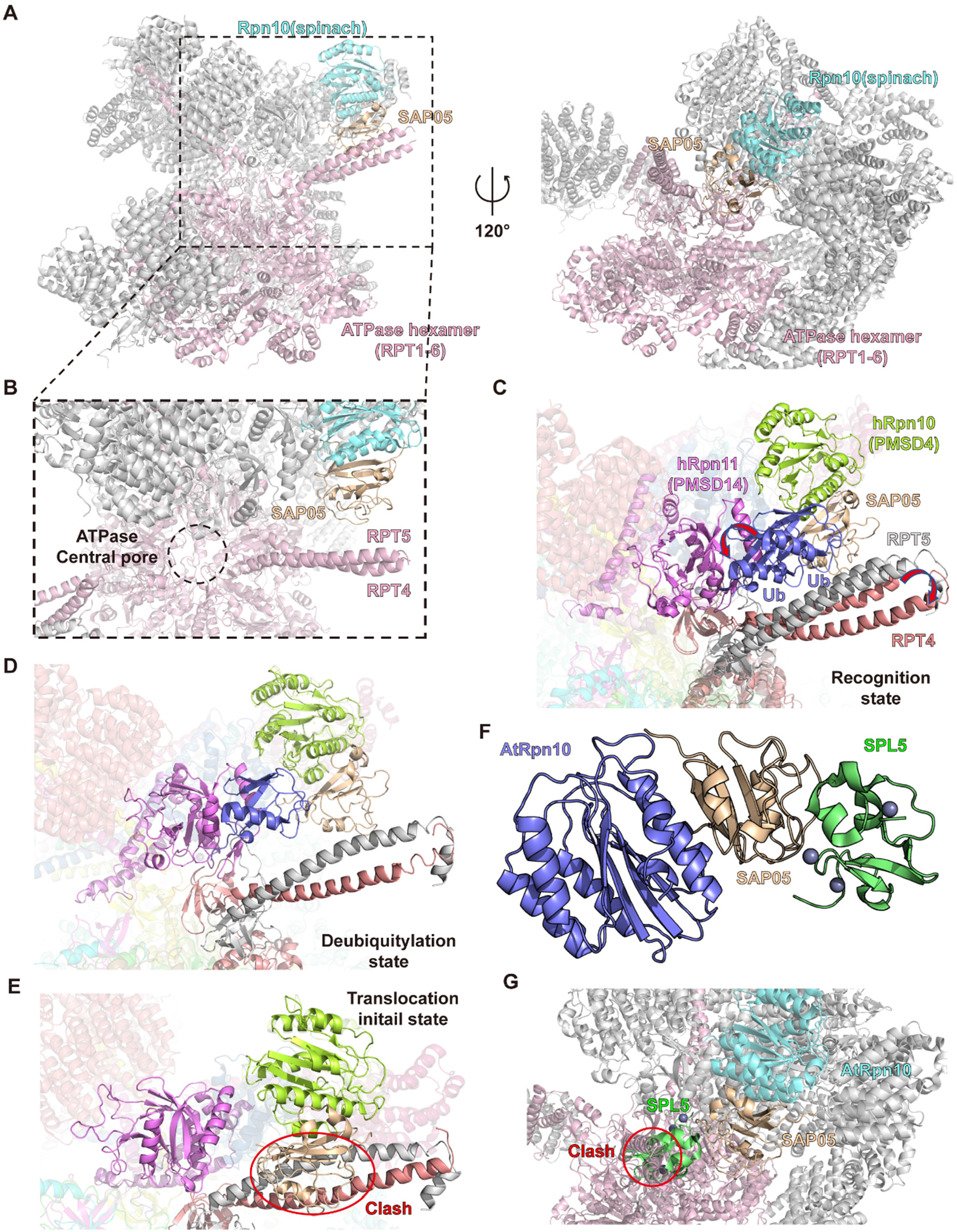
Superposing models of SAP05-Rpn10 complex in the proteasome. A) B) Different views of superposing models of SAP05 biding with spinach proteasome 19S regulatory particle (PDB: 8AMZ). C) D) E) Superpose our AYWB_SAP05-AtRpn10 structure to the different states of substrate-engaged human 26S proteasome: recognition state (PDB: 6MSB, 6MSD, the conformation difference between these two structures are pointed out by arrows), deubiquitylation state (PDB: 6MSE) and translocation initial state (PDB: 6MSH, the clash is marked by red circle). F) Superpose the crystal structure of AYWB_SAP05-SPL5_ZnF complex (PDB: 8PFC) to our AYWB_SAP05-AtRpn10 structure. G) Superpose the AtRpn10-AYWB_SAP05-SPL5_ZnF ternary complex structure model to the spinach proteasome 19S regulatory particle.

A spatiotemporal continuum model of the breakdown of polyubiquitylated protein by human 26S proteasome has been resolved using cryo-EM by Dong *et al*. recently^18^. In the initial recognition state of the proteasome-substrate complex, there are two ubiquitins on the substrate docking on the N-terminal CC (coiled-coil) domain of RPT4–RPT5 next to RPN10 (Figure 5C). They will shift away from RPT4-RPT5 N-termini, Rpn10 and toward Rpn11 for deubiquitination during the substrate peptide inserts to the central pore (Figure 5D). The SAP05 is located close to these ubiquitin-binding regions and thus may work as the substitution of ubiquitin to bring substrate to the substrate entrance of the proteasome lid.

The SAP05 will not collide with any subunit in the 19S regulatory particle until the substrate translocation initiates. But once the substrate-engaged proteasome enters the translocation-initial state, a conformational rotation will occur in the proteasome lid, which will make the RPT4-RPT5 N-terminal CC domain occupy the binding site of SAP05 (Figure 5E). This clash suggests that the SAP05 is supposed to dissociate from Rpn10 before the translocation initiation.

In Sum, SAP05 is capable of bringing transcription factor proximal to the entrance of the central pore and the ubiquitin-binding region, which can initiate substrate degradation. And the SAP05 may leave from Rpn10 before translocation initiation to make the substrate unfolding process work smoothly.

## Discussion

According to previous *in vitro* degradation assays, ∼20% of cellular proteins are able to be degraded by proteasome independent of ubiquitination^19^. The plant proteasomal subunit Rpn10, which is reported to recognize both ubiquitinated molecules and UbInPD(ubiquitin-independent proteasomal degradation)-associated shuttling factors, can be hijacked by bacterial effector SAP05 to mediate ubiquitin-independent degradation of parasitifer’s transcription factor during phytoplasma invasion.

In this work, we resolved the crystal structure of two Rpn10-SAP05 complexes, and the authenticity of our structures was validated by the SPR binding affinity assay. Our crystal data revealed the recognition mechanism between SAP05 of two different species and plant Rpn10 in this process. These SAP05 adopt similar globular conformations containing a β-sheet core layer and three α-helices. They can dock to the same Rpn10 α1/α2 interface by the collaboration of the V-shape peptide and C-terminal region, mediated by a few conserved hydrogen bonds and hydrophobic interactions, and the conserved conformation of the SAP05 interface.

At the same time that we wrote this manuscript, Liu *et al*. reported the crystal structure of AYWB_SAP05-AtRpn10_vWA structure same as us and a structure of SAP05 interacting with its transcription factor target, SPL5_ZnF^20^. The SPL5_ZnF binds to the loop surface consisting of L5 and L6 loops in our AYWB_SAP05 structure (Figure 5f). After superimposing their SAP05-ZnF_SPL5 complex structure, our AYWB_SAP05-AtRpn10 structure and the structure of spinach proteasome 19S particle, the SPL5 will be located between SAP05 and RPT4-RPT5 CC domain (Figure 5g). However, the location of SPL5 has clashes with the RPT4-RPT5, which indicates the interaction between SPL5 and ATPase subunits and a conformation change during SPL5 binding.

In conclusion, our work reveals a conserved recognizing mode between phytoplasma pathogenic protein SAP05 and plant proteasomal subunit Rpn10 during the phytoplasma invasion. Besides the process of SAP05 hijacking SPL and GATA transcription factor families, another phytoplasma protein, SAP54, was reported to utilize Rad23 to mediate proteasomal degradation of MADS-box TFs. In consideration of the severe economic loss caused by phytoplasma diseases of crops, these hijack processes are worth studying to facilitate phytoplasma prevention. In addition, a heterobifunctional tool of targeted degradation via direct 26S proteasome PSMD2 subunit recruitment is reported this year^14^. The SAP05/SAP54^10-12^ function of connecting host protein to the Rpn10/Rad23 to mediate ubiquitin/UBL-independent degradation can enlighten the development of similar human proteasome-targeting chimera molecules.

## Methods

### Plasmids

DNA sequence of hRpn10vWA(1-193aa) was cloned from youbio PSMD4 (NM_002810) cDNA and inserted to pGEX-6p-1 vector by ClonExpress II One Step Cloning Kit (Vazyme). Other genes were synthesized and operated codon optimization for expression in E. coli by GenScript. These genes were prefixed with HRV 3C cleavage sites and cloned into the corresponding vectors between the NdeI and XhoI sites. The pet 28a (+) vector was selected for SAP05 and At_Rpn10vWA used the pGEX-6p-1 vector.

### Protein Expression and Purification

All plasmids were expressed in BL21(DE3) Chemically Competent Cell (Transgene). The antibiotic plate corresponding to the resistance gene on the vector was used to screen out the E. coli colonies importing the target gene, and the monoclonal colonies were picked and cultured in 5 mL of LB (containing 1‰ antibiotics) at 37°C overnight. The bacterial solution was transferred to 1L LB (containing 1‰ antibiotics), and the culture was expanded in a shaker at 220rpm 37°C for 4-5h, until the OD600 reached 0.6-0.8. Cool the medium down to 16°C, add a final concentration of 0.4 mM IPTG into LB to induce expression of the target gene in E. coli, then adjust the shaker speed to 180 rpm and continue culture for 14-16h.

Collect bacterial strain expressing His6-SAP05 by centrifugation and suspended in Lysis Buffer1 (20 mM HEPES pH 7.5, 150 mM NaCl, 5% glycerol, 20 mM imidazole). After sonication and ultracentrifugation, cell lysates were incubated with Ni-NTA Beads 6FF at 4°C for 1 hour. Beads were washed with Lysis buffer until the G250 dye did not turn blue, then changed to high salt buffer (20 mM HEPES pH 7.5, 500 mM NaCl for 3-5 column volumes, after which they were reset to low salt buffer. The target proteins were eluted from the beads with 300 mM imidazole buffer (20 mM HEPES pH 7.5, 150 mM NaCl, 5% glycerol, 300 mM imidazole), concentrated, and further purified by Superdex 75 Increase column (Cytiva) on a ÄKTA pure system to isolate the protein in monomeric form.

GST-Rpn10vWA was purified by glutathione affinity chromatography using Lysis buffer 2 (20 mM HEPES pH 7.5, 150 mM NaCl, 5% glycerol, 2 mM DTT) and eluted with Lysis Buffer 2 GSH buffer containing 30 mM reduced glutathione. The elution was submitted to size exclusion chromatography using Superdex 200 Increase 10/300 GL (Cytiva). The high-purity fractions of protein samples were collected after being characterized by Coomassie-stained SDS-PAGE. Protein mutants were purified by the same method.

To gain the SAP05-Rpn10vWA complex used for crystal, two types of protein elution were mixed at a molar ratio of 1.2 to 1 according to concentrations determined via a Nanodrop. After adding one percent HRV 3C protease, protein solutions were dialyzed to enzyme digestion buffer(20 mM HEPES pH 7.5, 150 mM NaCl, 5% glycerol, 2 mM DTT) overnight. Protein solutions were incubated with glutathione beads to remove the GST tag, and the beads were washed with dialysis buffer until the G250 dye did not turn blue. Flow-though together with washing solution was diluted with Buffer A (20 mM HEPES pH 8.5) and loaded onto anion exchanger Source 15Q chromatography and eluted at around 14 mS/cm conductance by gradient against Buffer B (20 mM HEPES pH 8.5, 1 M NaCl). Peak fractions eluted at around 14 mS/cm conductance were collected and characterized by 12.5% SDS-PAGE (epizyme PAGE Gel Fast Preparation Kit) for the removal of GST-mixed complexes. Then the sample was concentrated and loaded onto Superdex 75 Increase 10/300 GL or HiLoad 16/600 Superdex 75 pg(Cytiva) equilibrated with SPR buffer (20 mM HEPES pH 7.5, 150 mM NaCl). Conduct SDS–PAGE analysis to determine the fraction of protein complex with high purity, and concentrated to about 20 mg/ml to obtain the sample for crystallization.

### Crystallization, data collection and processing

20 mg/mL AYWB_SAP05-AtRpn10 or OY_SAP05-AtRpn10 complex was used for crystal initial screening with sitting-drop vapor diffusion at 291K. Each drop contained 0.5 μL protein complex solution and 0.5 μL crystallization reservoir buffer. Satisfying crystals of the AYWB_SAP05-AtRpn10 complex grew under a reservoir solution containing 0.2 M Sodium malonate pH 5.0, 20% w/v Polyethylene glycol 3,350, while the crystals of OY_SAP05-AtRpn10 complex grew under 0.1M MES pH 6.5, 15% w/v PEG 6000, 5% v/v MPD. Crystals were cryo-protected in the solution with 17 % glycerol in the respective reservoir solution.

Both the diffraction of the AYWB_SAP05-AtRpn10 and OY_SAP05-AtRpn10 complex crystal was collected under the temperature of 100 K and wavelength of 0.979183 Å at the SSRF BEAMLINE BL10U2 of Shanghai Synchrotron Radiation Facility. Raw data was auto-processed and scaled with Aimless 0.7.7^21^. We used the Alphafold2-predicted model of AYWB_SAP05 (AF-Q2NK94-F1), OY_SAP05 (AF-Q6YQ57-F1), and AtRpn10 (AF-P55034-F1) for molecular replacement phasing in PHENIX 1.19.2_4158^22^. The model building was visualized and operated by COOT^23^. Crystallographic data collection and refinement statistics are presented in Table 1.

**Table 1.** Crystallographic data collection and refinement statistics.

|  | AYWB_SAP05-AtRpn10<sup>†</sup> (PDB 8JTK) | OY_SAP05-AtRpn10<sup>†</sup> (PDB 8JTL) |
| --- | --- | --- |
| **Data collection** |  |  |
| Space group | P 3<sub>1</sub> 2 1 | P 1 2 1 |
| Cell dimensions |  |  |
| *a*, *b*, *c* (Å) | 49.24, 49.24, 215.154 | 52.428, 39.386, 121.952 |
| $\alpha$, $\beta$, $\gamma$ (°) | 90.0, 90.0, 120.0 | 90.0, 93.781, 90.0 |
| Resolution range (Å) | 71.72 – 1.57 (1.65 – 1.57)<sup>a</sup> | 31.08 – 1.69 (1.73 – 1.69)<sup>a</sup> |
| No. of unique reflections | 42242 | 55704 |
| *R*<sub>merge</sub> | 0.056 (0.768)<sup>a</sup> | 0.169 (1.773)<sup>a</sup> |
| *R*<sub>pim</sub> | 0.021 (0.314)<sup>a</sup> | 0.074 (0.763)<sup>a</sup> |
| *R*<sub>meas</sub> | 0.060 (0.831)<sup>a</sup> | 0.185 (1.936) |
| *I*/$\sigma$(*I*) | 18.0 (2.6)<sup>a</sup> | 5.0 (1.0)<sup>a</sup> |
| *CC*<sub>1/2</sub> | 0.999 | 0.827 |
| Completeness (%) | 95.9 (100)<sup>a</sup> | 99.2 (98.6)<sup>a</sup> |
| Redundancy | 8.2 (6.9)<sup>a</sup> | 6.4 (6.4)<sup>a</sup> |
| **Refinement** |  |  |
| Resolution (Å) | 24.93 – 1.57 | 25.35 – 1.78 |
| No. reflections | 42142 | 47533 |
| *R*<sub>work</sub> / *R*<sub>free</sub> (%) | 18.70 / 21.54 | 18.40 / 21.85 |
| No. of non-hydrogen atoms | 2522 | 5062 |
| Protein | 2322 | 4652 |
| Ligand/ion | 0 | 0 |
| Water | 200 | 410 |
| *B* factors (Å<sup>2</sup>) | 38.0 (Average) | 35.0 (Average) |
| Protein<sup>b</sup> | 33<sup>b</sup> | 31<sup>b</sup> |
| Solvent | 44.7 | 38.4 |
| R.m.s. deviations |  |  |
| Bond lengths (Å) | 0.008 | 0.014 |
| Bond angles (°) | 1.033 | 1.356 |
| Clashscore | 4.0 | 10.0 |

<sup>a</sup> Values in parentheses are for the highest-resolution shell.<sup>b</sup> The median values of the occupancy-weighted average B-factor per residue.<sup>†</sup> The data are collected from one crystal.

### Surface Plasmon Resonance

SPR assay was carried out on Biacore S200 (Cytiva). At first, the ligand was diluted to 5 μg/mL by 10 mM sodium acetate (pH 4.5) and conjugated to Series S Sensor Chip CM5 (Cytiva) using the amide couple kit until the response value approached to setting level. Then use ethanolamine to block the remaining reactive sites in the channel. To continue, the analytes are diluted by SPR Buffer into a series of concentration gradients and run through the ligand binding cell one by one for Kinetics/Affinity testing. Eventually, we used the Biacore S200 Evaluation System to fit the curve and calculate the dissociation equilibrium constant.

### Pull Down assay

50 μg of GST-AtRpn10/hRpn10 was incubated with glutathione resin pre-equilibrated with SPR buffer (20 mM HEPES pH 7.5, 150 mM NaCl) for 2 h at 16°C. Washing beads until the G250 dye did not turn blue. And the resin was incubated with 40 μg OY_SAP05 for 2 h. Take one-tenth beads to the loading buffer after washing the unbinding protein. Then the samples were boiled at 95°C for 15 min and run on 4-20% polyacrylamide gels (GenScript).

## Acknowledgment

Qingyun Zheng was supported by National Key R&D Program of China, (No. 2022 YFC3401501). We thank the Tsinghua University Branch of China National Protein Science Facility, Tsinghua University, Beijing, for the device support of crystal screening (mosquito, sptlabtech) and checking (XtaLAB FR-X, Rigaku) and the guidance about the crystallization experiment from Ming Li (National Protein Science Facility, Tsinghua University). The data collection was supported by the Shanghai Synchrotron Radiation Facility. We thank the Center of Pharmaceutical Technology, Tsinghua University for support of our binding assay and the experimental guidance from Ting Wang (enter of Pharmaceutical Technology, Tsinghua University).

## Data availability

Crystal structures of the AYWB_SAP05-AtRpn10 and OY_SAP05-AtRpn10 complex have been published in the Protein Data Bank (PDB ID code 8JTK and 8JTL respectively).

## References

1. Goldberg, A. L., Protein degradation and protection against misfolded or damaged proteins. Nature 2003, 426 (6968), 895–9.

2. Pohl, C.; Dikic, I., Cellular quality control by the ubiquitin-proteasome system and autophagy. Science 2019, 366 (6467), 818–822.

3. Chirnomas, D.; Hornberger, K. R.; Crews, C. M., Protein degraders enter the clinic - a new approach to cancer therapy. Nat Rev Clin Oncol 2023, 20 (4), 265–278.

4. Craxton, A.; Butterworth, M.; Harper, N.; Fairall, L.; Schwabe, J.; Ciechanover, A.; Cohen, G. M., NOXA, a sensor of proteasome integrity, is degraded by 26S proteasomes by an ubiquitin-independent pathway that is blocked by MCL-1. Cell Death Differ 2012, 19 (9), 1424–34.

5. Orlowski, M.; Wilk, S., Ubiquitin-independent proteolytic functions of the proteasome. Arch Biochem Biophys 2003, 415 (1), 1–5.

6. Sheaff, R. J.; Singer, J. D.; Swanger, J.; Smitherman, M.; Roberts, J. M.; Clurman, B. E., Proteasomal turnover of p21Cip1 does not require p21Cip1 ubiquitination. Mol Cell 2000, 5 (2), 403–10.

7. Tsvetkov, P.; Reuven, N.; Shaul, Y., Ubiquitin-independent p53 proteasomal degradation. Cell Death Differ 2010, 17 (1), 103–8.

8. Makaros, Y.; Raiff, A.; Timms, R. T.; Wagh, A. R.; Gueta, M. I.; Bekturova, A.; Guez-Haddad, J.; Brodsky, S.; Opatowsky, Y.; Glickman, M. H.; Elledge, S. J.; Koren, I., Ubiquitin-independent proteasomal degradation driven by C-degron pathways. Mol Cell 2023, 83 (11), 1921–1935 e7.

9. Huang, W.; MacLean, A. M.; Sugio, A.; Maqbool, A.; Busscher, M.; Cho, S. T.; Kamoun, S.; Kuo, C. H.; Immink, R. G. H.; Hogenhout, S. A., Parasitic modulation of host development by ubiquitin-independent protein degradation. Cell 2021, 184 (20), 5201–5214 e12.

10. Kitazawa, Y.; Iwabuchi, N.; Maejima, K.; Sasano, M.; Matsumoto, O.; Koinuma, H.; Tokuda, R.; Suzuki, M.; Oshima, K.; Namba, S.; Yamaji, Y., A phytoplasma effector acts as a ubiquitin-like mediator between floral MADS-box proteins and proteasome shuttle proteins. Plant Cell 2022, 34 (5), 1709–1723.

11. MacLean, A. M.; Orlovskis, Z.; Kowitwanich, K.; Zdziarska, A. M.; Angenent, G. C.; Immink, R. G.; Hogenhout, S. A., Phytoplasma effector SAP54 hijacks plant reproduction by degrading MADS-box proteins and promotes insect colonization in a RAD23-dependent manner. PLoS Biol 2014, 12 (4), e1001835.

12. MacLean, A. M.; Sugio, A.; Makarova, O. V.; Findlay, K. C.; Grieve, V. M.; Toth, R.; Nicolaisen, M.; Hogenhout, S. A., Phytoplasma effector SAP54 induces indeterminate leaf-like flower development in Arabidopsis plants. Plant Physiol 2011, 157 (2), 831–41.

13. Rumpler, F.; Gramzow, L.; Theissen, G.; Melzer, R., Did Convergent Protein Evolution Enable Phytoplasmas to Generate ‘Zombie Plants’? Trends Plant Sci 2015, 20 (12), 798–806.

14. Bashore, C.; Prakash, S.; Johnson, M. C.; Conrad, R. J.; Kekessie, I. A.; Scales, S. J.; Ishisoko, N.; Kleinheinz, T.; Liu, P. S.; Popovych, N.; Wecksler, A. T.; Zhou, L.; Tam, C.; Zilberleyb, I.; Srinivasan, R.; Blake, R. A.; Song, A.; Staben, S. T.; Zhang, Y.; Arnott, D.; Fairbrother, W. J.; Foster, S. A.; Wertz, I. E.; Ciferri, C.; Dueber, E. C., Targeted degradation via direct 26S proteasome recruitment. Nat Chem Biol 2023, 19 (1), 55–63.

15. Kandolf, S.; Grishkovskaya, I.; Belacic, K.; Bolhuis, D. L.; Amann, S.; Foster, B.; Imre, R.; Mechtler, K.; Schleiffer, A.; Tagare, H. D.; Zhong, E. D.; Meinhart, A.; Brown, N. G.; Haselbach, D., Cryo-EM structure of the plant 26S proteasome. Plant Commun 2022, 3 (3), 100310.

16. Marshall, R. S.; Li, F. Q.; Gemperline, D. C.; Book, A. J.; Vierstra, R. D., Autophagic Degradation of the 26S Proteasome Is Mediated by the Dual ATG8/Ubiquitin Receptor RPN10 in Arabidopsis. Molecular Cell 2015, 58 (6), 1053–1066.

17. Rani, N.; Aichem, A.; Schmidtke, G.; Kreft, S. G.; Groettrup, M., FAT10 and NUB1L bind to the VWA domain of Rpn10 and Rpn1 to enable proteasome-mediated proteolysis. Nat Commun 2012, 3, 749.

18. Dong, Y.; Zhang, S.; Wu, Z.; Li, X.; Wang, W. L.; Zhu, Y.; Stoilova-McPhie, S.; Lu, Y.; Finley, D.; Mao, Y., Cryo-EM structures and dynamics of substrate-engaged human 26S proteasome. Nature 2019, 565 (7737), 49–55.

19. Baugh, J. M.; Viktorova, E. G.; Pilipenko, E. V., Proteasomes Can Degrade a Significant Proportion of Cellular Proteins Independent of Ubiquitination. J Mol Biol 2009, 386 (3), 814–827.

20. Liu, Q.; Maqbool, A.; Mirkin, F. G.; Singh, Y.; Stevenson, C. E. M.; Lawson, D. M.; Kamoun, S.; Huang, W.; Hogenhout, S. A., Bimodular architecture of bacterial effector SAP05 drives ubiquitin-independent targeted protein degradation. 2023, 2023.06.19.545293.

21. Evans, P. R.; Murshudov, G. N., How good are my data and what is the resolution? Acta Crystallogr D 2013, 69, 1204–1214.

22. Liebschner, D.; Afonine, P. V.; Baker, M. L.; Bunkoczi, G.; Chen, V. B.; Croll, T. I.; Hintze, B.; Hung, L. W.; Jain, S.; McCoy, A. J.; Moriarty, N. W.; Oeffner, R. D.; Poon, B. K.; Prisant, M. G.; Read, R. J.; Richardson, J. S.; Richardson, D. C.; Sammito, M. D.; Sobolev, O. V.; Stockwell, D. H.; Terwilliger, T. C.; Urzhumtsev, A. G.; Videau, L. L.; Williams, C. J.; Adams, P. D., Macromolecular structure determination using X-rays, neutrons and electrons: recent developments in Phenix. Acta Crystallographica Section D-Structural Biology 2019, 75, 861–877.

23. Emsley, P.; Lohkamp, B.; Scott, W. G.; Cowtan, K., Features and development of Coot. Acta Crystallogr D 2010, 66, 486–501.

